# Magnetically powered chitosan milliwheels for rapid translation, barrier function rescue, and delivery of therapeutic proteins to the inflamed gut epithelium

**DOI:** 10.1101/2022.10.10.511568

**Authors:** M.J. Osmond, E. Korthals, C.J. Zimmermann, E.J. Roth, D.W.M. Marr, K.B. Neeves

## Abstract

Inflammatory bowel disease (IBD) is mediated by an overexpression of tumor necrosis factor-α (TNF) by mononuclear cells in the intestinal mucosa. Intravenous delivery of neutralizing anti-TNF antibodies can cause systemic immunosuppression and up to one-third of people are non-responsive to treatment. Oral delivery of anti-TNF could reduce adverse effects; however, it is hampered by antibody degradation in the harsh gut environment during transit and poor bioavailability. To overcome these shortcomings, we demonstrate magnetically powered hydrogel particles that roll along mucosal surfaces, provide protection from degradation, and sustain local release of anti-TNF. Iron oxide nanoparticles are embedded into a crosslinked chitosan hydrogel and sieved to produce 100-200 μm particles called milliwheels (m-wheels). Once loaded with anti-TNF, these m-wheels release 10% to 80% of their payload over one week at a rate that depends on crosslinking density and pH. A rotating magnetic field induces a torque on the m-wheels that results in rolling velocities greater than 500 μm/s on glass and mucus-secreting cells. The permeability of TNF challenged gut epithelial cell monolayers was rescued in the presence of anti-TNF carrying m-wheels which both neutralized the TNF and created an impermeable patch over leaky cell junctions. With the ability to translate over mucosal surfaces at high speed, provide sustained release directly to the inflamed epithelium, and provide barrier rescue, m-wheels demonstrate a potential strategy to deliver therapeutic proteins for the treatment of IBD.

## Introduction

Inflammatory bowel disease (IBD), which includes Chron’s disease (CD) and ulcerative colitis (UC), has an incidence in North America of 20.2 per 100,000 person years.^1^ While the precise cause is unknown, the leading hypothesis is that these are diseases of the gastrointestinal tract (GIT) that result from dysregulated immune responses to luminal antigens, such as bacteria, in genetically susceptible individuals.^2,3^ Associated with this disease, lesions in the small intestine and colon are characterized by their loss of barrier function, invasion of gut microbiota, and subsequent activation of the innate immune response. Part of this immune response includes the production of pro-inflammatory cytokines, such as tumor necrosis factor-α (TNF), which is secreted from mononuclear cells in the mucosa.^4^

Anti-TNF therapy is effective in inducing and maintaining remission, and in reducing hospitalization and intestinal resection.^5^ Yet, with widespread use over the last 20 years, several limitations persist; for example, up to one-third of patients are non-responders to anti-TNFs and an additional 10-20% lose response each year with treatment.^6^ Additionally, intravenous delivery of anti-TNFs can lead to side effects including infections, autoimmune reactions, and malignancy.^7^ In IBD, TNF is increased in the stool, but less so in the blood, suggesting a rationale for targeting the luminal side of the gut. Testing this approach, oral delivery of anti-TNFs is effective in animal models of IBD and has shown promise in phase I/II clinical trials.^8,9^ One example is the IgG antibody AVX-470, which attenuates degradation by nine-fold compared to human IgG antibodies.^10^ In addition, oral administration of AVX-470 inhibits gut inflammation in murine models of IBD,^11^ is well tolerated up to 3.5 g/day in humans,^9^ and yields biomarkers consistent with anti-TNF effects in the colon mucosa.^12^ While results are positive, only a small fraction of oral drugs reach lesions, requiring that human doses be on the scale of multiple grams per day, which is a costly and impractical amount to manufacture. This limitation underscores the necessity of protecting therapeutic proteins during oral delivery and the development of targeting approaches that can concentrate them at the site of inflammation.

To protect proteins from degradation in the GIT, hydrogels that reside in the stomach, or slowly transit through the GIT, can deliver therapeutics over prolonged periods.^13–16^ These hydrogels have successfully delivered insulin,^16,17^ contraceptives,^18^ and antiretrovirals^19^ into the bloodstream, but do not target a specific region of the GIT. Alternatively, microbots, micron-scale robots often powered by catalytic or enzymatic propulsion or external magnetic fields, can be swallowed, travel to the intestines, and release their payload in response to pH change, resident enzymes, or by remote actuation.^20–22^ Microbots designed to release a controlled amount of therapeutic often only do so over a period of hours rather than days due to payload limitations, which is suboptimal for treating chronic illness like IBD.^23^

In previous work, we have shown that magnetic microwheels (μ-wheels), disk-like assemblies of superparamagnetic polystyrene microparticles, can translate at speeds of 100’s μm/s along surfaces when driven by 1-10 mT rotating magnetic fields,^24–26^ deliver therapeutic proteins,^27,28^ and behave like swarms through complex vessels.^29^ Here we build upon that work to create hydrogel-based milliwheels (m-wheels) designed for translation along mucosal tissues and controlled release of therapeutic proteins with swarm-like behavior. These m-wheels are synthesized by embedding paramagnetic iron oxide nanoparticles in chitosan microparticles with sizes defined by sieve-based fabrication. We show that m-wheels roll at velocities of approximately 500 μm/s across glass and mucus producing cells and release anti-TNF antibodies for up to one week. In addition, m-wheels with or without anti-TNF loading were shown to rescue barrier function in an *in vitro* model of the TNF-inflamed gut epithelium.

## Materials and Methods

### Materials

Low molecular weight chitosan (50-190 kDa, cat#448869), glacial acetic acid (A6283), methotrexate (M9929-25MG), fluorescein isothiocyanate (FITC) isomer I (F7250-1G), Pluronic-F127 (P2443), MilliCell cell culture inserts (polycarbonate 0.4 μm pore size, PIHP01250), FITC-dextran 40,000 MW (53379), collagen from calf skin (C8919-20ML), TRI-reagent (T9424-25ML), and HT-29 cells (91072201-1VL) were purchased from Sigma Aldrich (Burlington, MA). (Tridecafluoro-1,1,2,2- tetrahydrooctyl)trichlorosilane (FOTS, SIT8174.0), was obtained from Gelest (Morrisville, PA). Iron oxide nanoparticles were obtained from Alpha Chemicals (black iron oxide, New Brighton, PA). Polydimethylsiloxane (PDMS, Sylgard 184 elastomer kit) was purchased from Krayden (Denver, CO). Colorectal adenocarcinoma cells (Caco-2) were purchased from ATCC (Manassas, VA). Fetal bovine serum (S11550) was obtained from Biotechne (Minneapolis, MN). High glucose-DMEM (HG-DMEM, 12-100-046), non-essential amino acids (11-140-050), sodium pyruvate (11360070) and penicillin/streptomycin (pen/strep. 10378016) solutions were purchased from Fisher Scientific (Waltham, MA). The High Capacity cDNA Reverse Transcription kit (4368814) and PowerUp SYBR Green Master Mix (A25741) were obtained from Applied Biosystems (Beverly Hills, CA). All primers were purchased from IDTDNA (Coralville, Iowa) with sequences provided in Table 1. Recombinant Alexa Fluor 555 anti-mucin 5AC antibody (ab218714), anti-occludin antibody (ab216327, and goat anti-rabbit IgG H&L (Alexa Fluor 488, ab150077) antibody were all obtained from Abcam (Waltham, MA). Anti-tumor necrosis factor-α (aTNF, 500-M26) and tissue necrosis factor-α (TNF, 300-01A) were purchased from Peprotech (Canbury, NJ). Magnets used for cell culture purchased from V&P Scientific (San Diego, CA) were 13 mm in diameter, 1.7 mm thick polytetrafluoroethylene (PTFE) coated and autoclavable (VP 779-13). All chemicals were used as received unless otherwise stated.

**Table 1.**
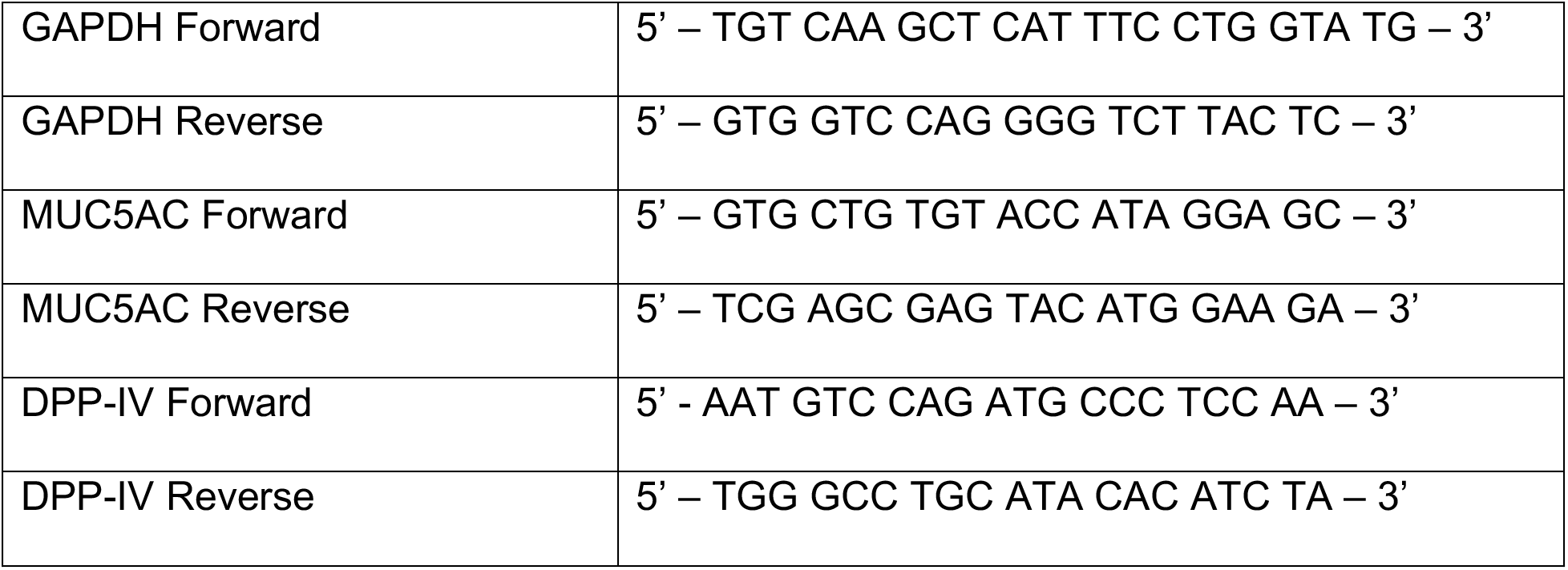
Sequences of primers used for the qPCR evaluation of GAPDH, MUC5AC, and DPP-IV.

### Synthesis and characterization of chitosan m-wheels

Magnetic chitosan m-wheels were synthesized using a sieving process (Fig. 1A).^30^ In this, low molecular weight chitosan was first purified by dissolving in 1% acetic acid to a concentration of 1% (w/v), vacuum filtered through a 22 μm filter, and then lyophilized. The purified chitosan was then dissolved in deionized H_2_O (diH_2_O) at 4% (w/v) along with 4% (w/v) iron oxide powder. This mixture was passed between two 30 mL syringes connected with a two-sided luer lock to produce a homogenous solution and mixed overnight on a rotator at room temperature to ensure full dissolution of the chitosan.

**Figure 1.**
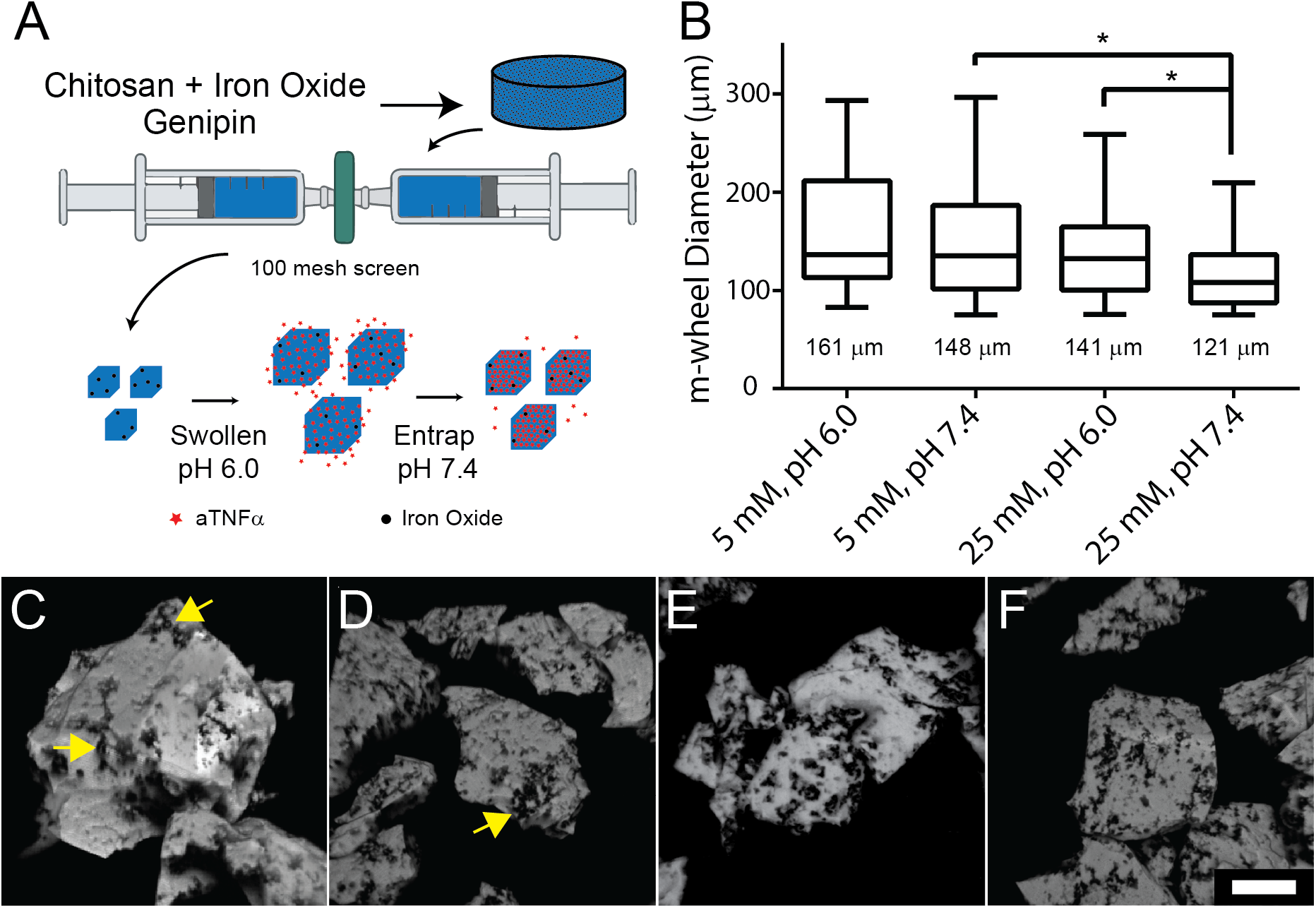
A. Schematic illustrating the fragmentation process for synthesizing magnetic chitosan m-wheels and loading of aTNF by swelling. B. Size distributions and mean diameters of the chitosan m-wheels as function of genipin concentration and pH, n = 200, * p<0.0001. Representative three-dimensional reconstructions from confocal microscopy images of iron oxide loaded chitosan m-wheels for synthesis conditions of 5mM pH 6.0 (C), 5 mM pH 7.4 (D), 25 mM pH 6.0 (E), 25 mM pH 7.4 (F). Scale bar = 50 μm. Dark inclusions within the particles are regions of iron oxide nanoparticles.

After dissolution was complete, 1 mL of the chitosan solution was transferred to a 3 mL syringe. To this, a separate 3 mL syringe containing 50 μL of genipin dissolved in absolute ethanol was attached using a luer lock connector, mixed for 30 s and then left in the syringe and rotated overnight in a humidity-controlled incubator. After curing, the gel was placed into a 10 mL syringe with 3 mL of water and mixed back and forth to another syringe of equal size to mechanically fragment the gel into smaller particles. These chitosan particles were then passed through a 100-mesh stainless steel screen (152 μm pore size, McMaster-Carr) five times and gravity filtered through a 200-mesh stainless steel screen (76 μm, McMaster-Carr) to remove the small particles. The particles retained on the screen are the m-wheels. m-Wheels used for experiments under sterile conditions were first submerged in 70% ethanol for 1 h under a UV lamp and then rinsed three times in sterile diH2O before storage in sterile buffer. All m-wheels were stored with approximately equal volumes of particles and buffer until use.

m-Wheel sizes were measured by fluorescence microscopy for gels made with 5 and 25 mM genipin and in solutions of pH 6.0 and pH 7.4 PBS buffer. m-Wheels were imaged between two coverslips with an Olympus IX81 using an Olympus UPlanFLN 4X (NA=0.13) objective, taking advantage of chitosan autofluorescence (ex/em 561/580 nm). Using Fiji, the particle edges were determined by thresholding each image using the *isoData* method to calculate the Feret diameter of each.^31^ Outliers for particle diameter identified via the ROUT method were removed from the data.^32^

### Caco-2 and HT-29-MTX cell culture

Caco-2 cells were cultured in HG-DMEM with 20% FBS and 1X pen/strep. HT-29 cells were cultured in HG-DMEM containing 10% FBS inactivated at 56°C for 1 h, 1X pen/strep, 1X sodium pyruvate and 1X non-essential amino acid. Cells were cultured at 37°C and 5% CO_2_. HT-29 cells were differentiated to a mucus-producing phenotype (HT-29/MTX) by culturing with media containing 10 μM methotrexate for 4 weeks with regular media changes every 2 days and passaged as needed.^33^ To confirm cell differentiation, qPCR was used to measure the relative expression of mucin (MUC5A) and dipeptidyl peptidase IV (DPP-IV). mRNA was isolated from HT-29 and HT-29/MTX differentiated cells using TRI-reagent and then converted to cDNA using the Applied Biosystems high-capacity cDNA reverse transcription kit. Relative expressions were measured using a StepOnePlus Real-Time PCR system from Applied Biosystems. The relative expression of MUC5A and DPP-IV was measured with GAPDH as a house keeping gene. MUC5A was measured to confirm the presence of mucin. DPP-IV is a known marker for enterocyte differentiation. Expression levels were evaluated using the ΔΔCt method.^34^

Cells were fixed with 10% neutral buffered formalin, permeabilized with 0.1% Triton X-100 for 15 min and stained by incubating in PBS containing recombinant Alexa Fluor 555 anti-mucin 5AC antibody 1/50 dilution. Confocal images were taken using an Olympus IX83 with a 40X (NA=0.60) objective equipped with a CSU-W1 confocal spinning unit from Yokogawa. Laser spectra used were 405 nm, 488 nm, and 561 nm. These images were used to verify the presence of MUC5AC on the surface of the cells.^33^ To produce a surface similar to those found in the native GIT, HT-29/MTX cells were co-cultured with Caco-2 cells on glass slides for rolling experiments. This coculture helps produce a more biomimetic ratio of cells, while preventing the bunching up of HT-29/MTX cells into a heterogenous layer. All co-culture experiments were seeded using a ratio of 9:1 Caco-2:HT-29/MTX as previously described.^35^ Glass slides for rolling experiments were seeded with 100,000 cells in 1 mL of media for the whole device and allowed to come to 90% confluence before rolling m-wheels on them.

### Milliwheel translation

In the presence of a rotating magnetic field and due to wet friction with available surfaces, m-wheels roll. We generate the necessary magnetic field using a custom apparatus^36^ in which two pairs of coils are mounted in the x and y plane with a fifth coil in the z direction above the sample (Fig. 2A). Appropriate currents are produced by an analog output card (NI-9263, National Instruments, Austin, TX), amplified (EP2000, Behringer, Willich, Germany), and sent to each pair of coils (Fig. 2B). The circular, rotating magnetic field is positioned normal to the rolling surface and the direction of the m-wheels manipulated by varying the ratio of current supplied to the x and y coils (Fig. 2C). The magnetic field strength and frequency are controlled with custom software.^37^

**Figure 2.**
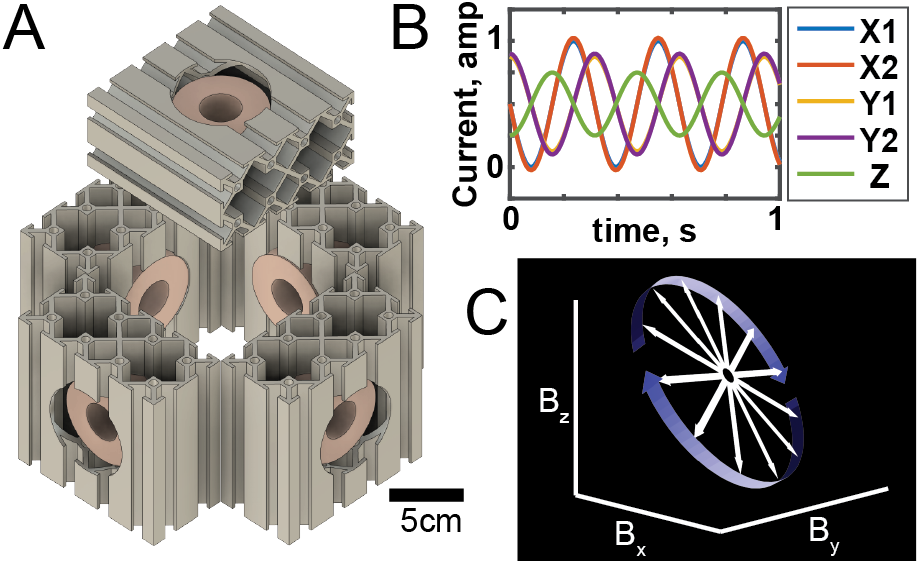
Magnetic apparatus constructed from five coils encapsulated by extruded aluminum housing contains two x-coordinate (X1 and X2), two y-coordinate (Y1 and Y2), and one z-coordinate coil (Z). B. Example waveforms amplified to produce an alternating magnetic field, one for each coil. Paired coils received the same signal to increase the magnetic field strength while providing uniform magnetic field gradient in the middle of the coil pair. C. Illustration of the time varying magnetic field vector (white) to create a rotational motion (blue).

To characterize rolling, a chamber was constructed by plasma bonding a PDMS chamber with an open top (30 mm in length, 8 mm width, 5 mm depth) to a glass slide. For experiments on glass, the slide was silanized with FOTS and blocked with 1% Pluronic F-127 in PBS for 15 min. For experiments on epithelial surfaces, Caco-2 and HT-29/MTX were co-cultured in the chamber as described above to 90% confluency. m-Wheels were collected using a magnet and all supernatant was removed, then resuspended in ~5X the volume of PBS buffer (rolling on glass) or DG-DMEM (rolling on cells), 50 μL of which was used within each chamber. m-Wheel rolling velocities were recorded using a Dino-lite Pro II digital microscope positioned under the sample to visualize the full channel width. Velocities were measured at two field strengths, 1.5 and 2.5 mT, and three frequencies, 5, 10, and 15 Hz using custom software.^36,38^

### Controlled release studies of aTNF

Chitosan m-wheels were collected via magnet while decanting and then swelled with addition of 5 mL of a pH 6.0 PBS solution. An aTNF antibody was fluorescently labeled by combining a 50 μL solution of aTNF (10 mg/mL) in carbonate buffer (pH 9.0) with a 2 μL solution of FITC in DMSO (10 mg/mL) for 10 h to form a bond between the isothiocyanate from FITC and a primary amine from aTNF. The products were then diluted with PBS to 0.67 mg/mL and purified using a Zeba Desalt Spin Column. FITC-labeled aTNF (FITC-aTNF) was added to the swollen m-wheels to a concentration of 10 μg/mL and allowed to load overnight at 37°C. After loading, the m-wheels were separated with a magnet, the solution collected to measure loading efficiency, and then a pH 7.4 PBS solution added to contract the m-wheels for 2 h. These m-wheels were then washed by collecting with a magnet and replacing with PBS three times before initiating release.

To measure release, each m-wheel solution was diluted 2X with PBS and 400 μL (200 μL of chitosan m-wheels) was pipetted into the apical side of each cell culture insert. To initiate release, 600 μL of either pH 6.0 or pH 7.4 PBS was added to the basolateral side of the cell culture insert. Each plate was then covered and placed in an incubator at 37°C and 95% relative humidity. At each time point, 600 μL was removed from each well, stored, and replaced with fresh PBS. Release was measured using a BioTek Synergy 2 plate reader quantifying FITC fluorescence and compared to a standard curve (ex/em 480/520 nm).

### Epithelial barrier permeability

Caco-2 cells were monocultured on cell culture inserts for permeability measurements. Inserts were seeded with 50,000 cells and grown for one week with 400 μL of media on the apical side and 600 μL on the basolateral. After one week, cells were transitioned to air-liquid interface culture (ALI) where only the basolateral media remained. Media was changed every two days until day 14 when it was changed daily. All inserts were treated with 50 ng/mL TNF to simulate an inflammatory environment. The inserts were treated with four different conditions, m-wheels that contain aTNF, m-wheels without aTNF, a positive control of media containing aTNF (0.5 μg/mL), and a negative control with neither m-wheels nor aTNF. The inserts treated with m-wheels were inoculated with 100 μL of the chitosan m-wheels loaded with ~1 μg of aTNF. An array of magnets was placed on the bottom of the well plate to generate a 10 mT magnetic field to hold the m-wheels onto the cell layer. In a separate set of experiments the magnets were excluded. After 48 h of treatment, the media from both sides of the membrane was removed. PBS (600 μL) was placed on the basolateral side and a 100 μg/mL FITC-conjugated-dextran in PBS (400 μL) was placed on the apical side of the membrane. After 4 h, the media from the basolateral side was collected and stored. The relative fluorescence values were measured using a plate reader (ex/em 480/520 nm). The relative permeability *P* of each cell layer was calculated using

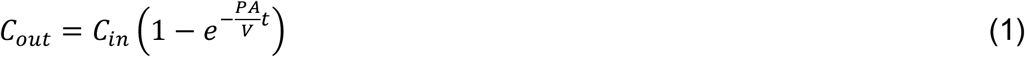

where *C_in_* is the initial concentration of FITC-dextran (100 μg/ml) in the apical side and is assumed to not change significantly over the course of the experiment, C_out_ is the concentration in the basolateral side, *A* is the area of the permeable membrane (0.6 cm^2^), *V* is the volume of the basolateral side (0.6 mL), and *t* is time.^39^ The concentration of the FITC-dextran on the apical side was measured at the conclusion of the experiment to confirm that *C_in_* does not significantly change.

### Confocal microscopy of TNF treated Caco-2 cells cultured on cell inserts

After cells were cultured and treated above for permeability studies, two of the six cell membranes were then rinsed with PBS and fixed in 10% neutral buffered formalin for 15 min. Cells were then rinsed with PBS and permeabilized using 0.1% Triton X-100 for 15 min. Cells were then labeled with anti-occludin antibody overnight at 4°C at 1/100 dilution and rinsed with PBS followed by goat anti-rabbit IgG H&L (Alexa Fluor 488) secondary antibody at 1/1000 dilution for 1 h. Just before imaging by spinning disk confocal microscopy, cells were incubated with 1 μg/mL DAPI for 15 min.

### Statistical Analysis

All distributions of m-wheels were tested for normality using a Shapiro-Wilks normality test and were found to be a non-normal distribution. To evaluate the variance between particle sets, a non-parametric Kruskal-Wallis test was used to measure the analysis of variance. If the Kruskal-Wallis test indicates a value of p < 0.05, this means a significant difference exists and each set was compared individually using a Dunn’s multiple comparisons test. P-values are reported in the figures.

## Results

### Characterization of chitosan m-wheels size and shape

m-Wheels were fabricated by crosslinking a chitosan solution containing iron oxide nanoparticles with genipin in bulk and mechanical fragmentation through a 152 μm sieve (Fig. 1A). The particle size distribution was measured as a function of the cross-linker genipin concentration and pH (Fig. 1B). A decrease in pH from 7.4 to 6.0 had no significant effect on particle size for 5 mM genipin crosslinking but did cause a decrease in particle size for 25 mM genipin crosslinking, indicating that swelling is dependent on crosslinking concentration as previously shown.^30,40,41^ m-Wheels crosslinked with 5 mM genipin are larger than m-wheels crosslinked with 25 mM genipin at pH 7.4. Chitosan autofluorescence (ex/em 561/580 nm) was used to visualize the shape of the m-wheels and the iron oxide distribution within them (Fig. 1C-F). These data show that the sieve-based fabrication method can create polygonal particles sized 100-200 μm that incorporate iron oxide nanoparticles.

### Mucus producing HT-29 cell differentiation to model the gut epithelium

HT-29 cells differentiated with methotrexate were characterized by measuring the relative mRNA expression of both MUC5AC and DPP-IV using qPCR. DPP-IV expression, a marker for a mucus producing phenotype in HT-29 cells, significantly increased by four-fold after differentiation (Fig. 3A).^42^ Both HT-29 and HT-29/MTX cells expressed MUC5AC, the major constituent of mucin found in the GIT, but not at a significantly different amount (Fig. 3A). To confirm mucin excretion from the cells, HT-29/MTX cells were fixed and stained for MUC5AC. Fig. 3B is a representative image of HT-29/MTX cells with the mucin visible in the cell interstitial space demonstrating that HT-29/MTX cells provide a mucus-rich surface for testing m-wheel translation. By coculturing HT-29/MTX cells with Caco-2 cells, a monolayer of cells surface similar to that of the GIT is achieved.^43,44^

**Figure 3.**
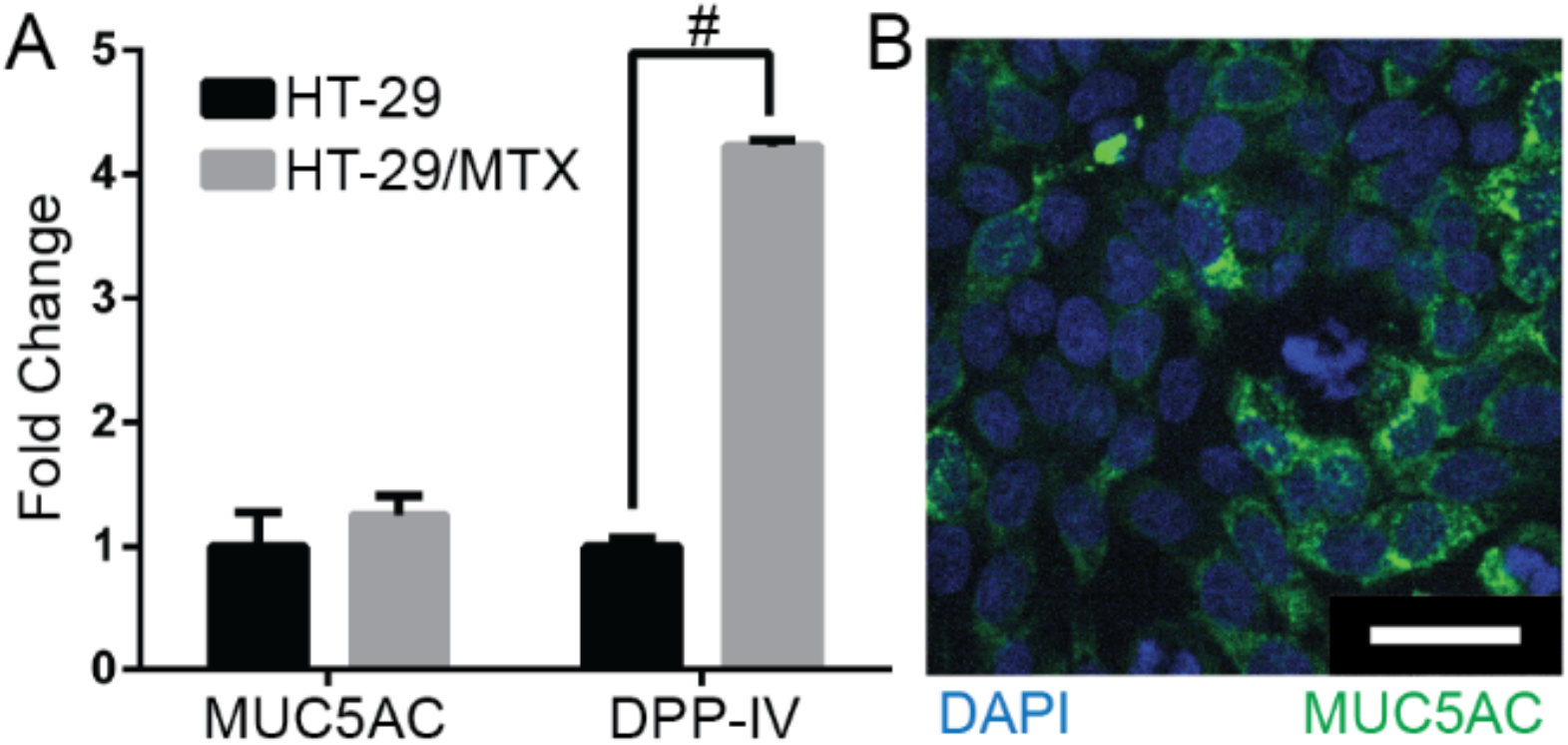
A. qPCR quantification of mucin (MUC5AC) and dipeptidyl peptidase-4 (DDP-IV) with undifferentiated HT-29 cells used as the control, # p < 0.05. B. Confocal micrograph of co-cultured Caco-2 and HT-29/MTX cells. Blue = cell nuclei (DAPI); Green = anti-MUC5AC. Scale bar = 50 μm.

### Milliwheel translation on glass and mucus producing co-culture

m-Wheel translation velocity was measured on glass or co-cultured Caco-2 and HT-29/MTX cells as a function of field frequency and field strength (Fig. 4, Supplemental Video 1). Co-cultured cells were chosen to provide a cellular topology and mucus rich surface to mimic the GIT cells. From these measurements, it is apparent that larger m-wheels travel faster than smaller m-wheels for a given magnetic field condition reflecting m-wheels with a larger circumference travel a greater distance per rotation. This observation is in agreement with our previous reports using micron-scale superparamagnetic polystyrene beads.^24^ The non-normal distribution of the velocities reflects a small number of large particles with very high translation velocities. There is also an increase in the mean velocity for m-wheels translating on glass as the field frequency increases (Fig. 4A). Because the rolling velocity is not linearly proportional to the field frequency these data indicate that the m-wheels are not rolling in phase with the magnetic field rotation. Here, the torque due to viscous drag and surrounding fluid exceeds that of the torque supplied by the magnetic field, in agreement with our previous observations at high frequencies with smaller particles.^24^

**Figure 4.**
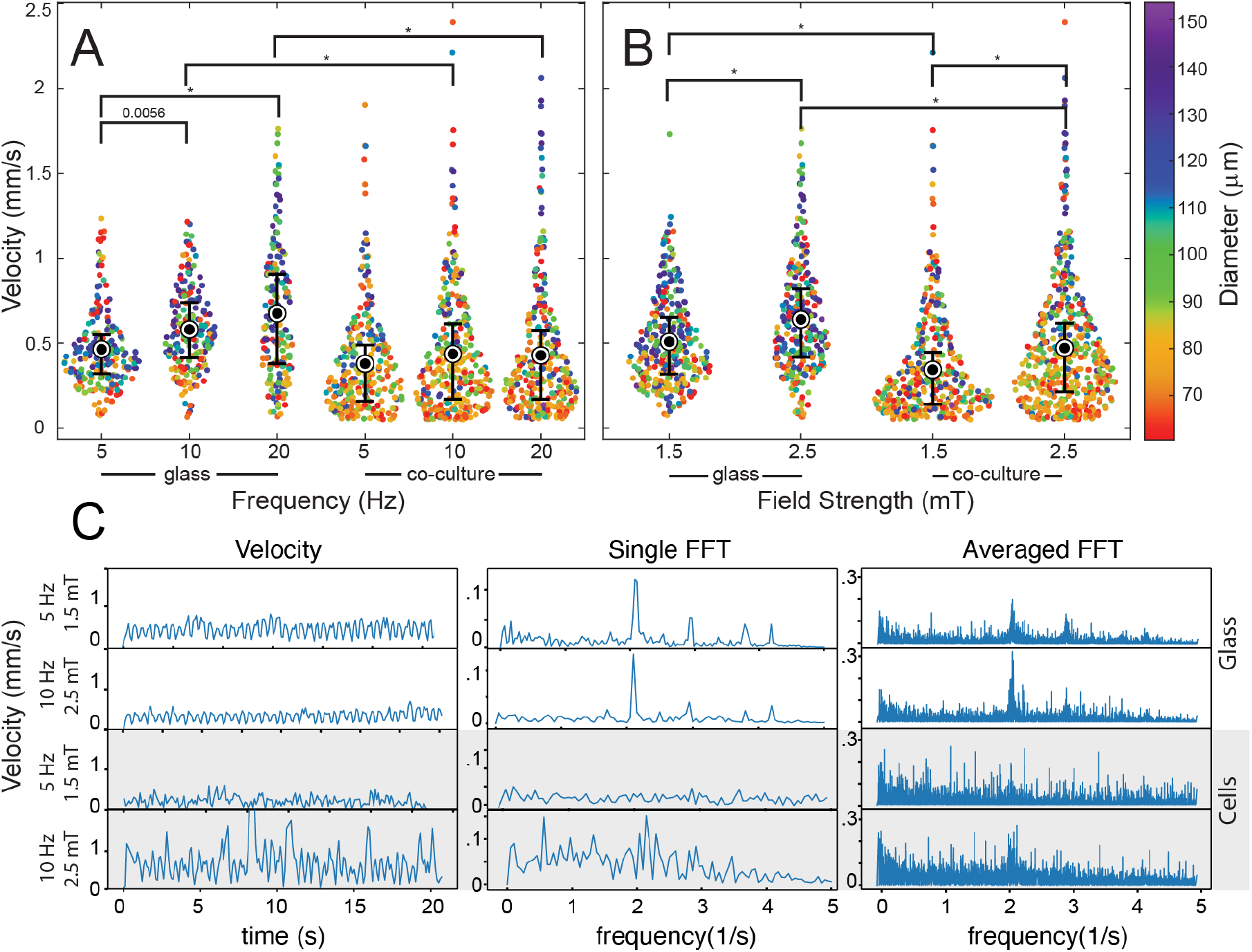
Velocity distributions of m-wheels translating on either glass or co-cultured surfaces as a function of magnetic field frequency (A) and magnetic field strength (B). Colors denote particle diameter. Black circles are the mean of the velocity, and the error bars are the 95% confidence intervals. N = 182 - 319 for frequency and N = 268 - 494 for field strength. P-value expressed on the figure, or * if p <0.0001. (C). Instantaneous velocity of a representative particles, fast Fourier transform of the same particles, and the combined fast Fourier transform of all particles (n = 94, 89, 199, and 204, top to bottom) under the same conditions. Each particle or set of particles labeled with corresponding frequency (5 or 10 Hz) and magnetic strength (1.5 or 2.5 mT), left axis, and they were rolled on glass (white background) or cells (gray background).

As expected, increasing the magnetic field strength from 1.5 mT to 2.5 mT results in an increase in translation velocity on both surfaces as more torque is created (Fig. 4B). m-Wheels roll slower on co-cultured cell monolayers than glass however, which appears to be a result of an aperiodic rolling motion of individual m-wheels on cells compared to glass. This is apparent in Fig. 4C where the instantaneous velocities and corresponding Fourier transforms of representative particles on glass show strong periodicity while those on cells do not. This periodicity, due to interactions with the surface and a combination of particle rotation rate and asymmetric particle features, is also apparent in Fourier transforms generated by averaging multiple particles. We note also no significant change in the translation velocity on cells across this range of frequencies, suggesting additional interactions not present on glass.

### Release of aTNF from m-wheels

The release rate of FITC-aTNF from m-wheels was measured for seven days as a function of crosslinking density (genipin concentration) and the pH of the suspending medium (Fig. 6). For a given pH, m-wheels with a low crosslinking concentration release aTNF faster than highly crosslinked gels. m-Wheels in pH 6.0 media released their payload faster than the pH 7.4 media due to chitosan swelling in accordance with m-wheel size measurements (Fig. 1B). While most of the curves follow a gradual release after day one, m-wheels cross-linked with 25 mM genipin and suspended in pH 6.0 media show a significant jump in release between three and seven days. This is a repeatable trend and suggests a faster breakdown of the chitosan network under this condition after three days.

**Figure 6.**
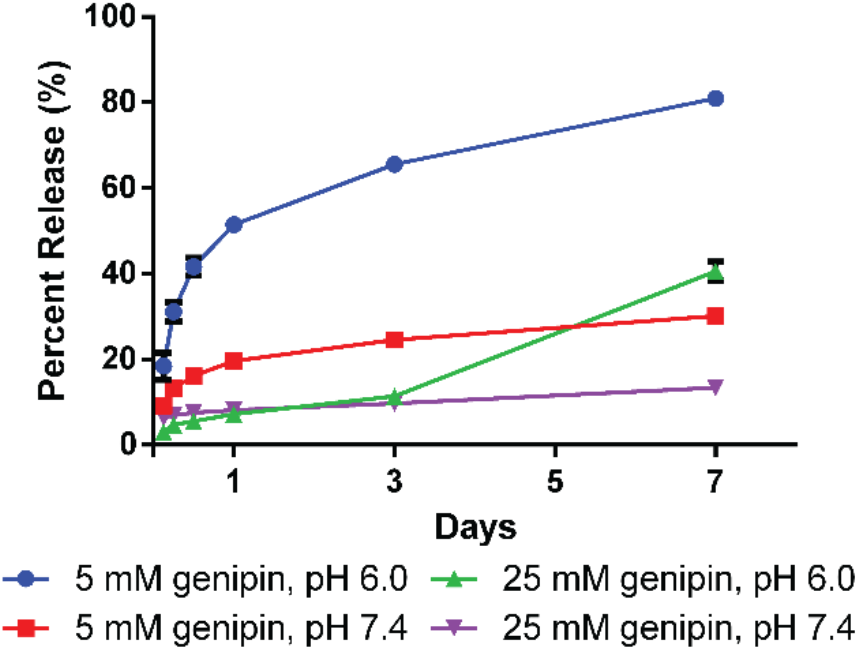
Release profile of FITC-aTNF from chitosan m-wheels cross-linked with either 5 mM or 25 mM genipin and released into PBS with pH of 6.0 or 7.4. Data presented as mean with standard deviation of n = 4. Lines connecting data points show trends for each condition.

### Milliwheels rescue barrier function of TNF treated gut epithelial cells

The intestinal epithelium functions as a semipermeable structure which allows nutrients to pass, while blocking pathogens like bacteria. To measure the ability of m-wheels to rescue or retard the degradation of this barrier function, Caco-2 intestinal cells were grown on cell culture inserts for two weeks as a model of the gut epithelium.^35,43,45^ To mimic an inflamed intestinal barrier, these cells were incubated with 50 ng/mL TNF for 48 h on the apical side of the membrane with four concurrent treatments: m-wheels, m-wheels loaded with aTNF, no treatment (negative control), and aTNF in solution (positive control). Each of these conditions was performed in the presence or absence of a magnetic field (~10 mT). Following the 48 h treatments, FITC-dextran was placed in the apical side of the membrane and the amount diffused through the cell layer was measured for 4 h. Using the change in concentration over time, the permeability was calculated for each condition using Eq. 1 (Fig. 7A). The higher the permeability measured, the lower the barrier function as induced by the TNF challenge.

**Figure 7.**
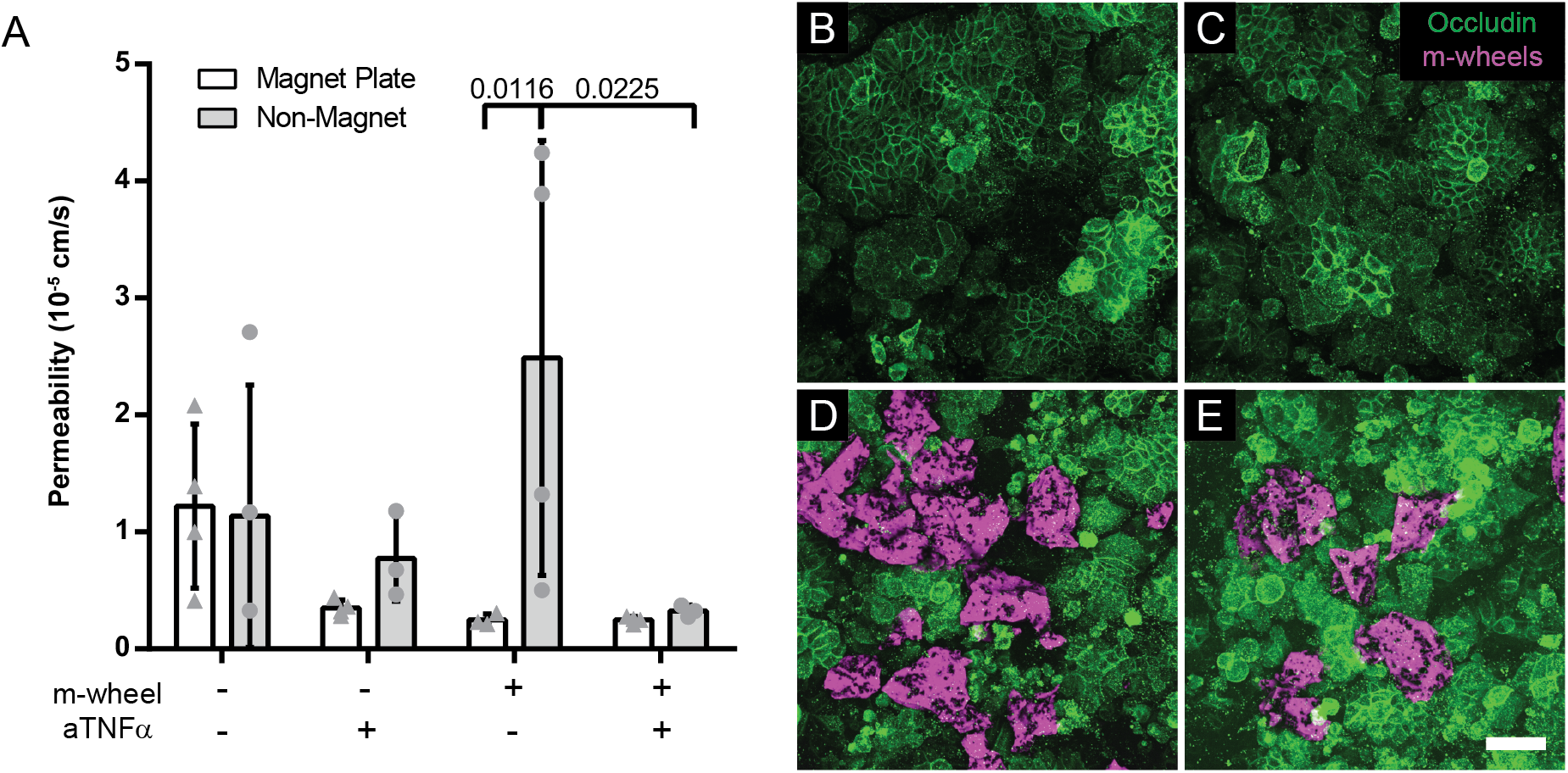
A. Permeability of the Caco-2 cells cultured for two weeks on cell culture membranes and treated with 50 μg/mL TNF for 48 h. Media with 100 μg/mL FITC-dextran was put in the apical side of the membrane and diffusion through the cell layer was measured after 4 h. Cells were cultured with (circles) and without (triangles) the presence of a magnetic field. B-E. Confocal micrographs from Caco-2 cells grown on cell culture inserts for 2 weeks; no treatment (B), aTNF only (C), m-wheels only (D), m-wheels and aTNFa treatment (E). Green = anti-occludin; magenta = chitosan autofluorescence. Scale bar = 50 μm. All images from inserts cultured in the presence of a magnetic field.

In the absence of a magnetic field, aTNF m-wheels showed a similar reduction in permeability as the positive control (Fig. 7A). m-Wheels without aTNF and no magnetic field did not rescue barrier function following the TNF challenge. The total amount of aTNF administered in the positive control and loaded in the m-wheels was equivalent (~100 ng). These data indicate that aTNF was biologically active when released from the chitosan and released at a rate that neutralizes TNF as well as a bolus of soluble aTNF.

In the presence of a magnetic field, m-wheels both with and without aTNF in the epithelial cell layer had permeabilities comparable to the positive control (Fig. 7A). These data suggest that the particles themselves can contribute to barrier function even in the absence of the neutralizing antibody. This trend is supported by images showing chitosan particles adhered to a significant fraction of the cell layer surface (Fig. 7D, E).

## Discussion

In an effort to create a hydrogel system for directed and controlled release of an anti-inflammatory drug to any location in the GIT, we have demonstrated a magnetically-manipulated, controlled-release microgel system that can travel over a mucus-rich model of the GIT lumen, release antibody over a period of a week, and exhibit rescue of epithelial barrier function in vitro. This was accomplished using a fabrication process based on sieving that can produce large numbers of microparticles between 100 and 200 μm. To avoid aggregation of iron oxide nanoparticles in emulsion based techniques, we fabricated bulk gels that are then broken down into smaller homogenous particles.^46^

m-Wheels translate at high speeds over both glass and mucus producing cell layers. At an average speed of 500 μm/s, these m-wheels could roll the 3.5 m distance of the ileum in just under 2 hours. Increasing the torque on the m-wheels allows them to break from the weak interactions between the chitosan and glass or cellular surfaces. Even though the chitosan is mucoadhesive and can stick to the mucus producing cells, enough torque is provided by the magnetic field to dislodge them for continued rolling. In this, m-wheels travel as swarms aiding delivery of high concentrations to a specific location. When the magnetic field is turned off, this mucoadhesive property holds the wheels in place while the antibodies are released.

m-Wheels were fabricated using chitosan to provide pH responsive particles to load and release an antibody payload. Swelling and encapsulation allows for increased loading efficiency without crosslinking the antibodies to the matrix.^30,47^ m-Wheels fabricated with different crosslinking densities and the pH of their respective release mediums alter the aTNF release. While the stomach has a pH of 1.5 to 3.0, which often requires capsules and other delivery coatings to protect therapeutic agents from degradation,^50,51^ the pH of the lower GIT fluctuates between 5.7 to 7.4.^48,49^ As such, we measured aTNF release at pH 6.0 and 7.4. The rate of aTNF released depends on the crosslinking density of m-wheels, with higher crosslinking slowing release. aTNF was released the fastest at pH 6.0 from the increased swelling of the chitosan. From this, the release of aTNF can be tuned to release an optimal amount depending on location or severity of the intestinal lesion.

Caco-2 cells were grown on cell membranes treated with TNF and then measured for a change in their permeability and morphology. The conditions treated with m-wheels either with or without release of aTNF exhibit a similar reduction in permeability as those treated with aTNF alone. We hypothesize that m-wheels decrease the permeability, in part, by adding a diffusive barrier, while also neutralizing TNF similar to the native mucus layer in healthy epithelium.^52,53^ Without a magnet, m-wheels are washed away and no longer act as a barrier for the damaged monolayer.

## Conclusions

To devise a potential treatment for people with chronic intestinal diseases such as IBD, we used well characterized and available hydrogels to make large quantities of magnetic m-wheels that can rapidly roll over mucosal surfaces. These m-wheels can be loaded with drugs such as aTNF at various concentrations and are designed to release from hours to weeks at the site of treatment. They provide a potential drug delivery system for targeted delivery of proteins that can be remotely guided to rapidly translate to inflamed portions the GIT.

## Acknowledgments

The authors acknowledge support from the National Institutes of Health under grants R21AI138214 and R01NS102465 as well as the University of Colorado School of Medicine’s GI and Liver Innate Immunity Program (GALIIP).

